# Rps3/uS3 promotes mRNA binding at the 40S ribosome entry channel and stabilizes preinitiation complexes at start codons

**DOI:** 10.1101/098699

**Authors:** Jinsheng Dong, Colin Echeverría Aitken, Anil Thakur, Byung-Sik Shin, Jon R. Lorsch, Alan G. Hinnebusch

## Abstract

The eukaryotic 43S pre-initiation complex (PIC) bearing Met-tRNA_i_^Met^ in a ternary complex (TC) with eIF2-GTP scans the mRNA leader for an AUG codon in favorable “Kozak” context. AUG recognition provokes rearrangement from an open PIC conformation with TC bound in a state not fully engaged with the P site (“P_OUT_”) to a closed, arrested conformation with TC tightly bound in the “P_IN_” state. Yeast ribosomal protein Rps3/uS3 resides in the mRNA entry channel of the 40S subunit and contacts mRNA via conserved residues whose functional importance was unknown. We show that substitutions of these residues reduce bulk translation initiation and diminish initiation at near-cognate UUG start codons in yeast mutants in which UUG selection is abnormally high (Sui^-^), conferring the Ssu^-^ phenotype. Two such Ssu^-^ substitutions—R116D and R117D—also increase discrimination against an AUG codon in suboptimal Kozak context. Consistently, the Arg116 and Arg117 substitutions destabilize TC binding to 48S PICs reconstituted in vitro with mRNA harboring a UUG start codon, indicating destabilization of the closed P_IN_ state with a UUG:anticodon mismatch. Using model mRNAs lacking contacts with either the mRNA entry or exit channels of the 40S subunit, we demonstrate that Arg116/Arg117 are crucial for stabilizing PIC:mRNA contacts at the entry channel, complementing the function of eIF3 at both entry and exit channels. The corresponding residues in bacterial uS3 promote the helicase activity of the elongating ribosome, suggesting that uS3 contacts with mRNA enhance multiple phases of translation across different domains of life.

## INTRODUCTION

Accurate identification of the translation initiation codon in mRNA by ribosomes is crucial for expression of the correct cellular proteins. This process generally occurs in eukaryotic cells by a scanning mechanism, wherein the small (40S) ribosomal subunit first recruits charged initiator tRNA (Met-tRNA_i_^Met^) in a ternary complex (TC) with eIF2-GTP, in a reaction stimulated by eukaryotic initiation factors (eIFs) 1, 1A, 3 and 5. The resulting 43S pre-initiation complex (PIC) attaches to the 5’ end of the mRNA and scans the 5’UTR with the TC bound in a metastable state, “P_OUT_”, suitable for inspecting successive triplets for complementarity with the anticodon of Met-tRNA_i_^Met^ in the P site, to identify the AUG start codon. Nucleotides surrounding the AUG, particularly at the −3 and +4 positions (the Kozak context), further influence the efficiency of start codon selection. In the scanning PIC, eIF2 can hydrolyze GTP, dependent on GTPase activating protein eIF5, but P_i_ release is blocked by eIF1, whose presence also impedes stable binding of Met-tRNA_i_^Met^ in the “P_IN_” state. Start-codon recognition triggers dissociation of eIF1 from the 40S subunit, allowing P_i_ release from eIF2-GDP·P_i_ and TC binding in the P_IN_ state of the 48S PIC (Fig. 1A). Subsequent dissociation of eIF2-GDP and other eIFs from the 48S PIC enables eIF5B-catalyzed subunit joining and formation of an 80S initiation complex with Met-tRNA_i_^Met^ base-paired to AUG in the P site (reviewed in (Hinnebusch 2014)).

**Figure 1.**
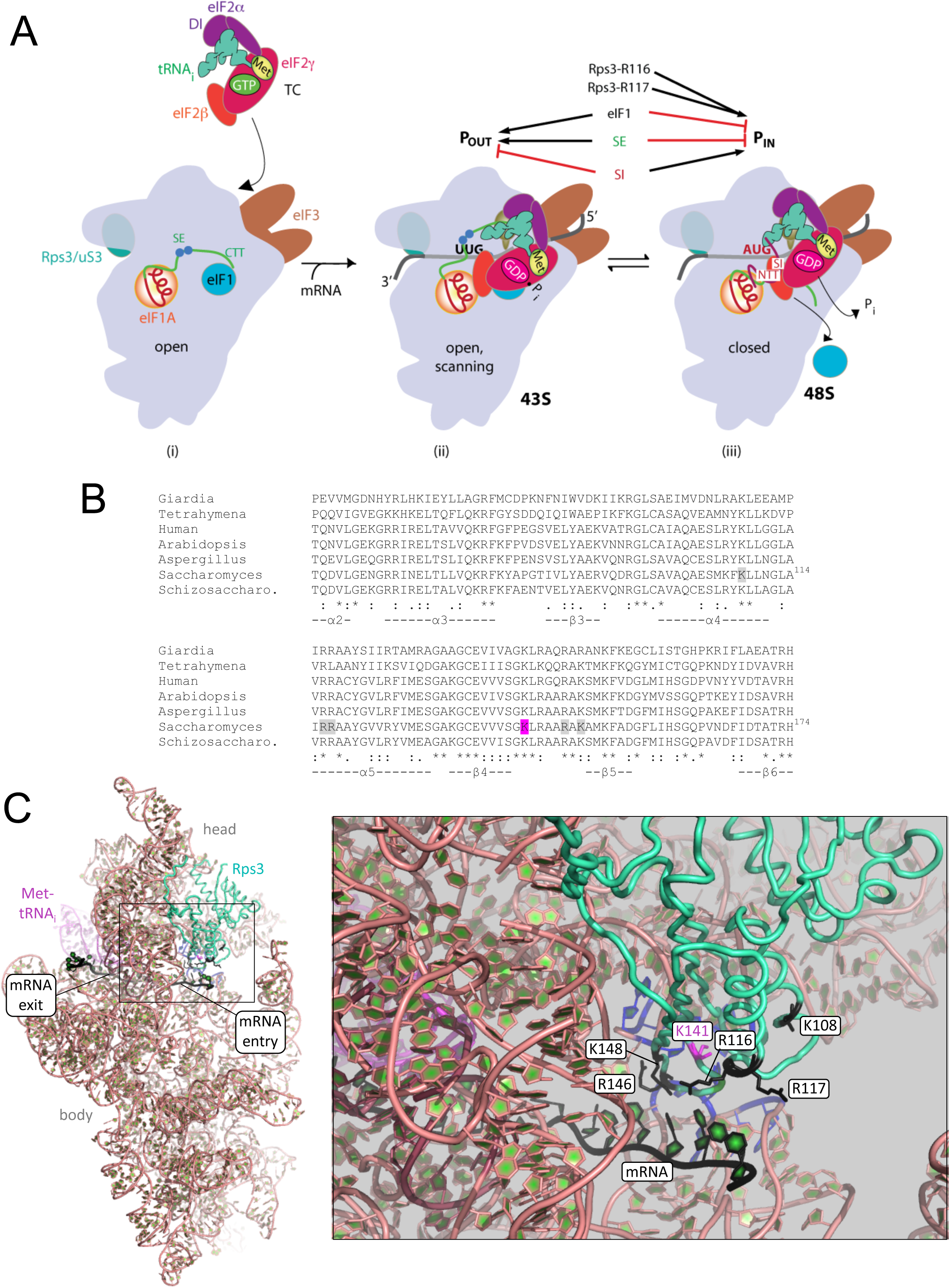
Rps3/uS3 plays a critical role in promoting mRNA binding at the 40S entry site and stabilizing the preinitiation complex at the start codon. (**A**) Model describing known conformational rearrangements of the PIC during scanning and start codon recognition. (i) eIF1 and the scanning enhancers (SEs) in the CTT of eIF1A stabilize an open conformation of the 40S subunit to which TC rapidly binds. Rps3 is located on the solvent exposed surface of the 40S near the entry channel; the bulk of eIF3 binds on the solvent-exposed surface with a prominent domain at the mRNA exit channel; (ii) The 43S PIC in the open conformation scans the mRNA for the start codon with Met-tRNA_i_^Met^ bound in the P_OUT_ state. eIF2 can hydrolyze GTP to GDP•P_i_, but release of P_i_ is blocked. (iii) On AUG recognition, Met-tRNA_i_^Met^ moves from the P_OUT_ to P_IN_ state, clashing with eIF1 and the CTT of eIF1A, provoking displacement of the eIF1A CTT from the P site, dissociation of eIF1 from the 40S subunit, and P_i_ release from eIF2. The NTT of eIF1A, harboring scanning inhibitor (SI) elements, adopts a defined conformation and interacts with the codon:anticodon helix. (Above) Arrows summarize that eIF1 and the eIF1A SE elements promote P_OUT_ and impede transition to P_IN_ state, whereas the scanning inhibitor (SI) element in the NTT of eIF1A stabilizes the P_IN_ state. Results presented below indicate that Rps3 residues R116/R117, in contact with mRNA at the entry channel, stabilize the P_IN_ state, and also promote PIC interaction with mRNA at the entry channel, complementing the role of eIF3 in PIC-mRNA interactions at the exit channel (Adapted from (Hinnebusch 2014)). (**B**) Alignment of Rps3 sequences from diverse eukaryotes. Six conserved residues of Rps3 at the mRNA entry channel analyzed in this study are highlighted in grey or Pink. (**C**) Position of Rps3 in the yeast 48S PIC, and locations of conserved Rps3 residues at the entry channel. The partial yeast 48S PIC (py48S; PDB3J81) is depicted on the left in cartoon format highlighting Rps3 (green), mRNA (black), Met-tRNA_i_^Met^ (Pink), and rRNA residues of h18 or h34 that comprise the entry channel latch (blue). The boxed region is amplified on the right where the six Rps3 residues analyzed here, which interact with mRNA or the rRNA latch, are highlighted in black or magenta, shown in stick format, and labeled. Other ribosomal proteins and eIFs 1, 1A, and 2 are hidden for clarity.

eIF1 plays a dual role in the scanning mechanism, promoting rapid TC loading in the P_OUT_ conformation while blocking rearrangement to P_IN_ at both near-cognate start codons (e.g. UUG) and cognate (AUG) codons in poor Kozak context; hence eIF1 must dissociate from the 40S subunit for start codon recognition (Fig. 1A). Consistent with this, structural analyses of partial PICs reveal that eIF1 and eIF1A promote rotation of the 40S head relative to the body (Lomakin and Steitz 2013) (Hussain et al. 2014a), thought to be instrumental in TC binding in the P_OUT_ conformation, but that eIF1 physically clashes with Met-tRNA_i_^Met^ in the P_IN_ state (Rabl et al. 2011; Lomakin and Steitz 2013), and is both deformed and displaced from its 40S location during the P_OUT_ to P_IN_ transition (Hussain et al. 2014a). Mutations that weaken eIF1 binding to the 40S subunit reduce the rate of TC loading and elevate initiation at near-cognate codons or AUGs in poor context as a result of destabilizing the open/P_OUT_ conformation and favoring rearrangement to the closed/P_IN_ state during scanning (Martin-Marcos et al. 2011; Martin-Marcos et al. 2013). Moreover, decreasing wild-type (WT) eIF1 abundance reduces initiation accuracy, whereas overexpressing eIF1 suppresses initiation at near cognates or AUGs in poor context (Valasek et al. 2004; Alone et al. 2008; Ivanov et al. 2010; Saini et al. 2010; Martin-Marcos et al. 2011). In fact, cells exploit the mechanistic link between eIF1 abundance and initiation accuracy to autoregulate eIF1 expression: the AUG codon of the eIF1 gene (*SUI1* in yeast) occurs in poor context and the frequency of its recognition is inversely related to eIF1 abundance (Ivanov et al. 2010; Martin-Marcos et al. 2011).

The stability of the codon-anticodon duplex is an important determinant of initiation accuracy, as the rate of the P_OUT_ to P_IN_ transition is accelerated and the P_IN_ state is stabilized in the presence of AUG versus non-AUG start codons (Kolitz et al. 2009). Favorable Kozak context might also contribute to P_IN_ stability (Pisarev et al. 2006; Martin-Marcos et al. 2011), but the stimulatory effect of optimum context on initiation rate is not well understood. It seems to require the α-subunit of eIF2 (Pisarev et al. 2006) and structural analyses of partial mammalian 43S (Hashem et al. 2013) and yeast 48S PICs (Hussain et al. 2014a) place the eIF2α domain-1 near the key -3 context nucleotide (nt) in the exit channel of the 40S subunit. The conserved β-hairPin of 40S protein uS7 (Rps5 in yeast) also occurs in this vicinity in a yeast partial 48S (py48S) PIC (Hussain et al. 2014b). We have shown that the β-hairPin of yeast Rps5 is important for both efficient and accurate translation initiation in vivo and the stability of P_IN_ complexes reconstituted in vitro (Visweswaraiah et al. 2015). Approximately seven additional mRNA nt upstream of the - 3 position occupy the 40S exit channel, and there is evidence that these 40S:mRNA interactions plus additional contacts between segments of eIF3 and mRNA nt protruding from the exit channel also enhance PIC assembly at the start codon (Kozak 1991; Pestova and Kolupaeva 2002; Pisarev et al. 2008; Aitken et al. 2016).

eIF3 is a multisubunit complex that binds directly to the 40S subunit and promotes both recruitment and stable association of TC and mRNA with the PIC, and enhances scanning and accurate start codon recognition in vivo (Valasek 2012). Recent structural analyses (Erzberger et al. 2014; Aylett et al. 2015; Llacer et al. 2015), combined with earlier biochemical and genetic studies (Valasek 2012), reveal that different subunits/domains of yeast eIF3 interact with the PIC at multiple sites, effectively encircling the PIC and interacting with both the mRNA-entry and - exit pores on the solvent-exposed surface, as well as with the decoding center on the interface surface of the 40S subunit. The major point of contact involves binding of a heterodimer of the PCI domains of the eIF3a and c subunits to the 40S solvent side, below the platform and exit channel pore. A second contact occurs near the entry channel and involves segments of the eIF3a, b, g, and i subunits; at least some of these interactions appear to be dynamic, as alternative contacts with eIFs anchored in the decoding center have also been observed (Erzberger et al. 2014; Aylett et al. 2015; Llacer et al. 2015). We recently presented evidence that the PCI domain of eIF3a mediates a key stabilizing interaction between the PIC and mRNA at the exit channel{Aitken, 2016 #7247}. Using a panel of m^7^G-capped, unstructured model mRNAs to reconstitute 48S PICs, we determined that the eIF3a PCI domain interaction at the exit channel is functionally redundant with mRNA:PIC interactions at the entry channel, and is essential for stable 48S PIC assembly when the mRNA is truncated in a way that leaves the entry channel largely empty. Other eIF3 domains/subunits contribute to the functionally redundant contacts at the entry channel, but are not essential even when the opposite exit channel is unoccupied by mRNA. This suggests that other components of the PIC, including elements of the ribosome itself participate in stabilizing mRNA binding at the entry channel.

In fact, the cryo-EM structure of a partial yeast 48S PIC revealed predicted contacts between mRNA 6-12 nt located 3’ of the AUG codon with particular amino acids of ribosomal protein Rps3/uS3 at the mRNA entry channel of the 40S subunit. Several Rps3 residues also appear to interact with 18S rRNA residues that form the “latch” on the entry channel, a non-covalent interaction between rRNA nt in helices 18 and 34 (Hussain et al. 2014b). Interestingly, while the latch is closed in the yeast 48S complexes with AUG in the P site and in crystal structures of other partial PICs (Rabl et al. 2011; Lomakin and Steitz 2013; Weisser et al. 2013), the latch is open in a recent cryo-EM structure of a PIC formed with mRNA containing an AUC codon, owing to upward movement of the head away from the body of the 40S subunit. The P-site is also widened in this open PIC conformation such that the tRNA_i_^Met^ is not fully engaged with rRNA residues in the body that contribute to the highly stable P_IN_ conformation observed in the corresponding AUG complex (Llacer et al. 2015). Hydroxyl radical probing of yeast PICs reconstituted with AUG or AUC mRNAs also revealed a more open conformation of the P site and less-constricted mRNA entry channel in the AUC complex (Zhang et al. 2015), consistent with a scanning-conducive conformation of the 40S subunit when a near-cognate triplet occupies the P site. Although Rps3 residues appear to interact with the mRNA and with rRNA residues of the entry channel latch, there is no functional evidence that these predicted contacts are important for the efficiency or fidelity of start codon recognition. Here we provide strong genetic and biochemical evidence that these Rps3 residues enhance the stability of the P_IN_ state and promote recognition of poor initiation sites in vivo, and also mediate stabilizing mRNA interactions with the PIC at the entry channel that are functionally redundant with eIF3-dependent PIC:mRNA interactions at the exit channel.

## RESULTS

### *RPS3* mutations restore discrimination against near-cognate UUG codons in a hypoaccurate eIF5 Sui^-^ mutant in vivo

To examine the role of Rps3 in the mRNA entry channel, we introduced single substitutions into six highly conserved basic residues found in proximity to either mRNA nt or rRNA residues in h18 or h34 of the entry channel latch in the cryo-EM structure of the partial yeast 48S PIC, with Met-tRNA_i_^Met^ base-paired with AUG in the P_IN_ state (Hussain et al. 2014b). These include Arg-116 and Arg-117 at the N-terminus of helix α5, and Lys-141, Arg-146, and Lys-148 in the loop between β-strands 4-5 (Figs. 1B-C). We also substituted Lys-108 (K108) at the C-terminus of α4, based on its proximity to the entry channel and predicted involvement, along with R116/R117 in ribosome helicase activity (Takyar et al. 2005; Graifer et al. 2014). Residues were substituted with Ala to shorten the side-chain, or with acidic residues to alter side-chain charge (Supplementary Table S1). The mutations were generated in an *RPS3* allele under its own promoter on a low-copy plasmid and examined in a yeast strain with wild-type (WT) chromosomal *RPS3* under a galactose-inducible promoter (*P_GAL1_*). Mutant phenotypes were scored following a switch from galactose to glucose, where *P_GAL1_-RPS3*^+^ expression is repressed. Mutations *K141A* and *R146A* were lethal and prevented growth on glucose medium, whereas others conferred slow-growth (Slg^-^) phenotypes that were very strong for *R116A*, moderate for *K141D*, and slight for *R146D* (Fig. 2A-B; summarized in Table S1).

**Figure 2.**
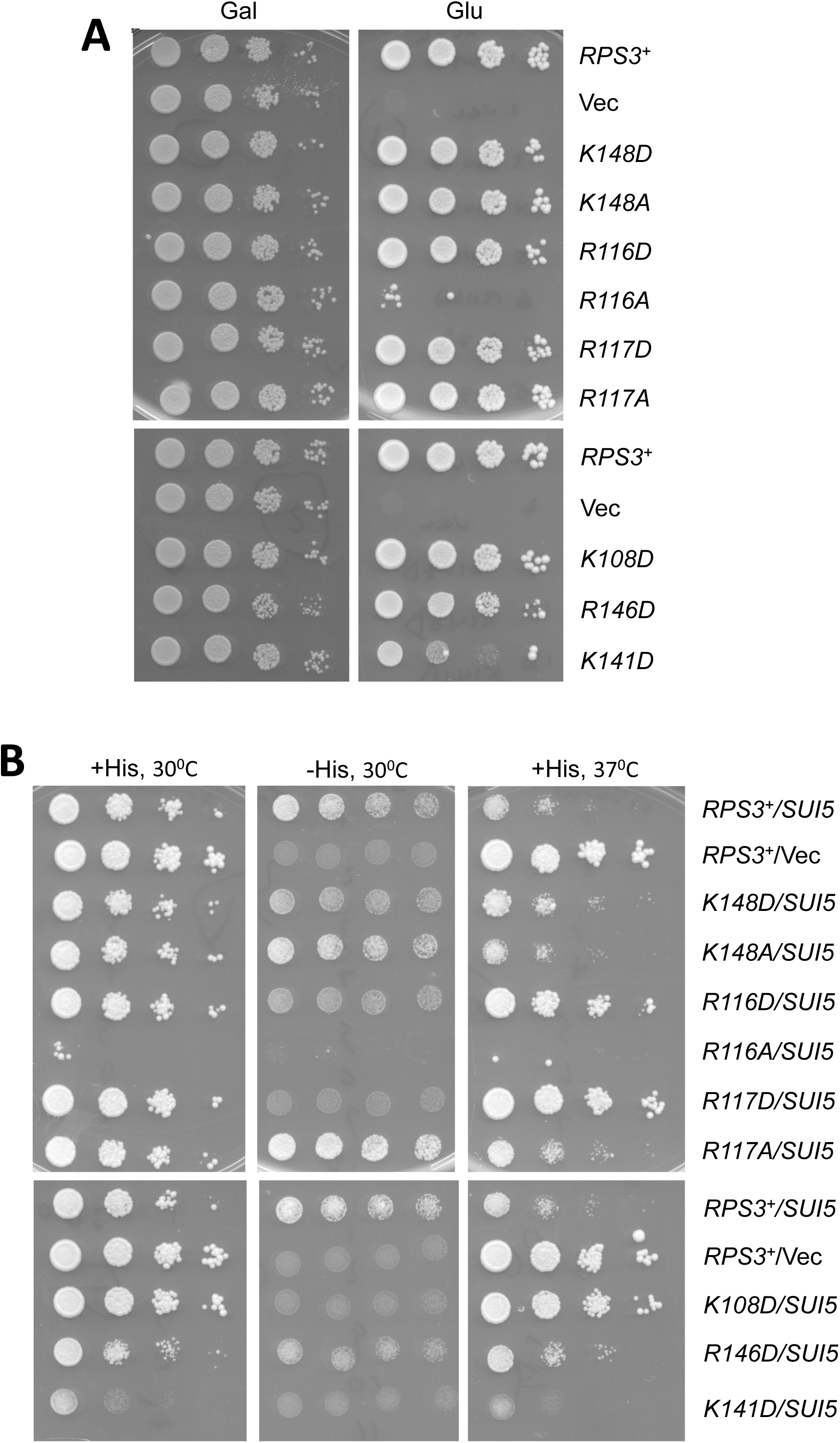
Certain *RPS3* alleles suppress both the His^+^ and Slg^-^ phenotypes conferred by eIF5 mutation *SUI5* in *his4-301* cells. (**A**) Certain *RPS3* alleles confer Slg^-^ phenotypes in otherwise WT cells. Serial dilutions of the following *P_GAL_-RPS3 his4-301* strain (HD2738) harboring the indicated plasmid-borne *RPS3* allele or empty vector (Vec) were spotted on S_GAL_-Leu or SC-Leu medium, and incubated for 3-4d at 30° C: *RPS3*^+^ (HD2754), Vector (HD2755), *K148D* (HD2765), *K148A* (HD2764), *R116D* (HD2767), *R116A* (HD2766), *R117D* (HD2769), *R117A* (HD2768), *K108D* (HD3120), *R146D* (HD2772), and *K141D* (HD2779). (**B**) Serial dilutions of the *his4-301* strains in (A) also containing sc *SUI5* plasmid p4281 or empty *TRP1* vector YCplac22 (Vec) were spotted on SC-L-W (+His plates) or SC-L-W-H supplemented with 0.0015 mM histidine (0.5% of the standard supplement; -His plates) and incubated at 30° C or 37° C for 3-5d.

To assess changes in initiation fidelity, the *RPS3* mutations were tested for the ability to suppress the histidine auxotrophy of *his4-301*, a mutant allele lacking the AUG start codon, by increasing initiation at the third, in-frame UUG codon and thereby restoring expression of the histidine biosynthetic enzyme His4. None of the *RPS3* mutations allowed detectable growth on glucose medium lacking histidine, suggesting that they do not reduce the stringency of selecting AUG as start codon. Accordingly, we tested them for the ability to suppress the elevated UUG initiation on *his4-301* mRNA and the resulting His^+^ phenotype conferred by the dominant Sui^-^ mutation *SUI5*, encoding the G31R variant of eIF5 (Huang et al. 1997). The His^+^ phenotype of plasmid-borne *SUI5* was diminished to varying extents by the *K148D, R116D, R117D, K108D*, and *R146D* alleles of *RPS3* (Fig. 2B, Table S1). (Although *K141D* also suppresses growth on −His medium, it has a similar effect on +His medium, unlike the other alleles just mentioned.) *SUI5* also confers a Slg^-^ phenotype in histidine-replete medium, particularly at elevated temperature (37°C), and this phenotype was also diminished strongly by *R116D, R117D,* and *K108D*, and to a lesser extent by *K148D* and *R146D* (Fig. 2B, Table S1). (Considering that *R146D* confers a Slg^-^ phenotype in otherwise WT cells (Fig. 2A), its ability to improve growth at 37 °C relative to WT *RPS3* in the *SUI5* background (Fig. 2B) indicates a greater ability to suppress the growth defect of *SUI5* than is evident from judging growth of *SUI5* strains alone.) These results suggest that a subset of Rps3 substitutions mitigate the effect of *SUI5* in reducing the accuracy of start codon recognition.

The effect of *SUI5* on the fidelity of start codon selection can be quantified by an increase in the expression of a *HIS4-lacZ* reporter containing a UUG start codon, as compared to the expression levels observed when the same cells are transformed with empty vector (Fig. 3A, columns 1-2). Supporting the interpretation that specific Rps3 substitutions dampen this effect, we found that all five *RPS3* mutations that reduce the His_+_/Sui^-^ phenotype of *SUI5* (Fig. 2B) also suppress the elevated expression of a *HIS4-lacZ* reporter containing a UUG start codon that is conferred by *SUI5* (Fig. 3A, col. 1 vs. 2-7). *SUI5* additionally produces a modest increase in expression of the matching reporter with an AUG initiation codon (Fig. 3B, cols. 1-2), an effect that is similarly reversed by the *RPS3* mutations (Fig. 3B, col. 1 vs. 3-7). Despite this last effect, all five *RPS3* mutations confer a marked reduction in the UUG:AUG initiation ratio for the two reporters (Fig. 3C, col. 1 vs. 3-7). These results demonstrate that the Rps3 substitutions restore the strong preference, typical of WT cells, for AUG versus UUG start codons in *SUI5* mutant cells, thus conferring strong Ssu^-^ phenotypes.

**Figure 3.**
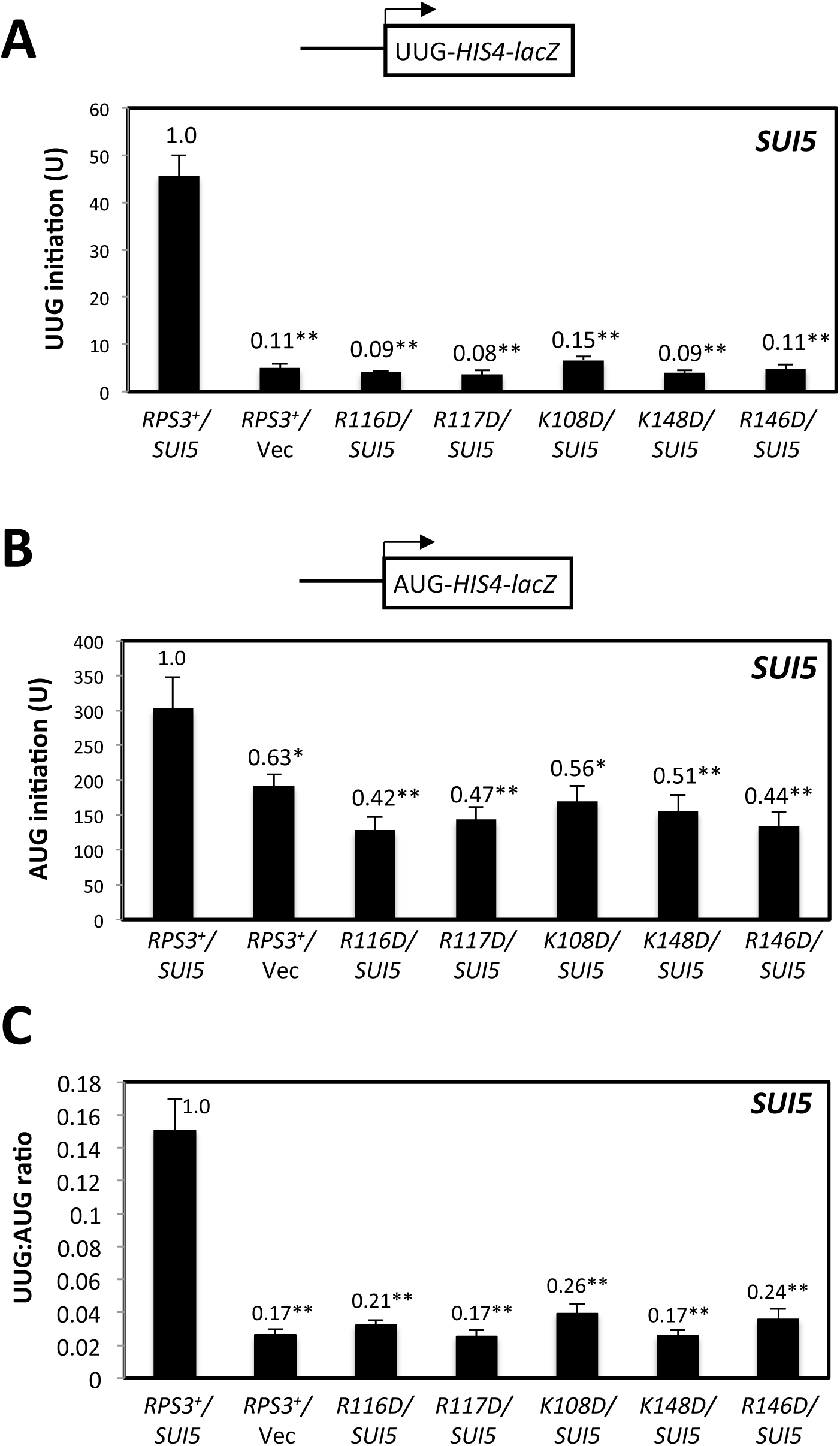
*RPS3* alleles suppress the elevated UUG:AUG initiation ratio of *HIS4-lacZ* reporters conferred by *SUI5*. (**A-C**) Transformants of *his4-301* strains with the indicated *RPS3* alleles and either *SUI5* plasmid p4281 or vector harboring the *HIS4-lacZ* reporters shown schematically with AUG (A) or UUG (B) start codons (plasmids p367 and p391, respectively) with AUG (A) or UUG (B) start codons were cultured in SD+His medium to an OD_600_ of 1.0-1.2, and β-galactosidase specific activities were measured in whole cell extracts (WCEs) in units of nanomoles of *o*-nitrophenyl-β-D-galactopyranoside (ONPG) cleaved per min per mg of total protein. Mean activities with S.E.M.s (shown as error bars) were determined from six independent transformants. In (C), mean ratios (with S.E.M.s) of expression of the UUG versus AUG reporter were calculated from the results in (A-B). Asterisks indicate statistically significant differences between each strain and the *RPS3/SUI5* strain determined by a two-tailed, unpaired Student’s t-test (*, P<0.05; **, P<0.01).

### *RPS3* Ssu^-^ mutations increase discrimination against the eIF1 AUG codon in suboptimal context

In addition to reducing initiation at near-cognate UUG codons in Sui^-^ mutants, previously identified Ssu^-^ substitutions in eIF1 and eIF1A were shown to intensify discrimination against the AUG start codon of the *SUI1* gene encoding eIF1, which occurs in poor Kozak context. This feature of *SUI1* initiation underlies negative autoregulation of eIF1 synthesis, dampening the ability to overexpress eIF1 in WT cells, as the excess eIF1 suppresses initiation at the *SUI1* start codon (Martin-Marcos et al. 2011). Consistent with these previous findings, four of the five *RPS3* Ssu^-^ substitutions reduce steady-state expression of eIF1, with the strongest reductions seen for *R116D* and *R117D*, and lesser effects for *R146D* and *K148D* (Fig. 4-B). *R116D* and *R117D* also decreased the expression of the WT *SUI1-lacZ* fusion containing the native, poor context of the *SUI1* AUG, but did not diminish expression of the *SUI1-opt-lacZ* reporter, in which the native *SUI1* context is replaced with an AUG in optimum context with A nucleotides at the upstream -1 to -3 positions (Martin-Marcos et al. 2011). Accordingly, *R116D* and *R117D* significantly increase the *SUI1-opt-lacZ/SUI1-lacZ* initiation ratio (Fig. 4C). Thus, in addition to suppressing UUG initiation, *R116D* and *R117D* clearly discriminate against the *SUI1* AUG codon in poor context. In contrast, *R146D* and *K148D* reduce eIF1 expression but have little effect on *SUI1-lacZ* expression, perhaps indicating that their ability to discriminate against the native *SUI1* initiation region, which is smaller in magnitude compared to *R116D/R117D* (Fig. 5A-B), requires other sequence or structural features not maintained in the *SUI1-lacZ* reporter. The fact that the *RPS3* Ssu^-^ mutants exhibit reduced expression of native eIF1 implies that their increased discrimination against the suboptimal context of the *SUI1* AUG codon overrides the expected effect of diminished eIF1 levels both in reducing discrimination against poor AUG context and boosting eIF1 synthesis, and of increasing, rather than decreasing, UUG initiation (Ivanov et al. 2010; Martin-Marcos et al. 2011).

**Figure 4.**
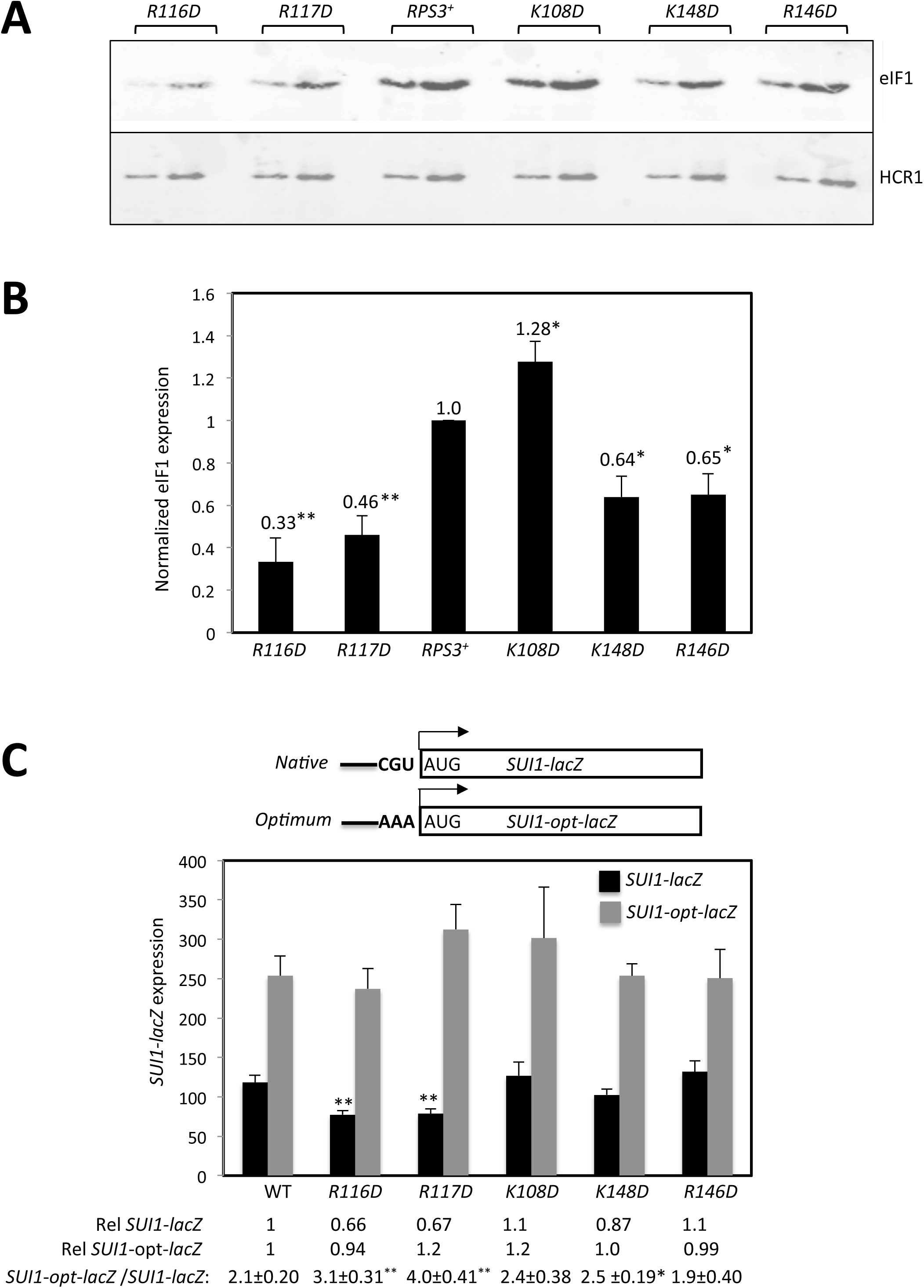
Certain *RPS3* Ssu^-^ alleles exacerbate discrimination against the AUG start codon of the *SUI1* gene encoding eIF1. (**A**) Strains described in Fig. 2A with the indicated *RPS3* alleles were cultured in SD+His+Trp+Ura medium to an OD_600_ of ~1.0 and WCEs were subjected to Western analysis using antibodies against eIF1 or Hcr1 (as loading control). Two amounts of each extract differing by a factor of two were loaded in successive lanes. (**B**) eIF1 expression, normalized to that of Hcr1, was obtained for each strain by quantifying the Western signals in (A), and mean values (±S.E.M.) were calculated from three biological replicates. Asterisks indicate significant differences between mutant and WT as judged by the Student’s t-test (P<0.005). (**C**) Strains from (A) also harboring *SUI1-lacZ* (pPMB24) and *SUIl-opt-lacZ* (pPMB25) reporters were cultured and assayed for β-galactosidase expression as described in Figure 3, except using SD+His+Trp medium. Mean expression levels and S.E.M.s from 6 transformants are plotted, and relative (Rel) mean expression levels normalized to that of the WT strain are listed below the histogram, along with the expression ratios for the *SUI1-lacZ* versus *SUI1-opt-lacZ* reporters. Asterisks indicate significant differences between mutant and WT as judged by a two-tailed, unpaired Student’s t-test (*, P<0.05; **, P<0.01).

**Figure 5.**
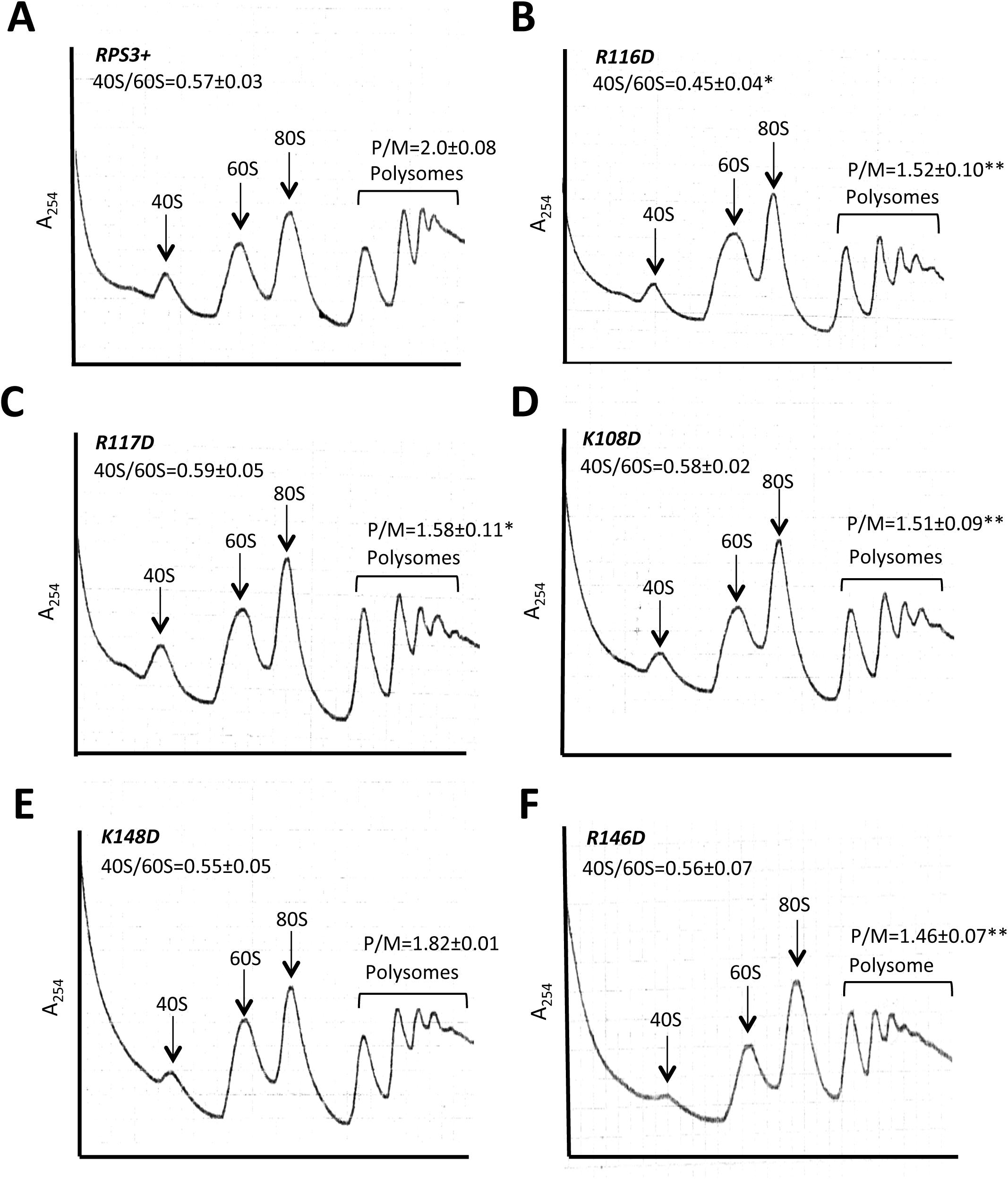
*RPS3* Ssu^-^ alleles reduce bulk translation initiation. Strains from Fig. 2A were cultured in SC-Leu at 30° C to A_600_ of ~1.0, and cycloheximide was added (50μg/ml) prior to harvesting. WCEs were separated by sucrose density gradient centrifugation and scanned at 254 nm. Mean (± S.E.M.) polysome/monosome (P/M) and free 40S/60S ratios from three biological replicates are indicated. Asterisks indicate significant differences between mutant and WT as judged by a two-tailed, unpaired Student’s t-test (*, P<0.05; **, P<0.01).

### *RPS3* Ssu^-^ mutations reduce bulk translation initiation without affecting 40S subunit abundance

Except for *K148D*, the other four *RPS3* Ssu^-^ mutations conferred moderate reductions in the ratio of polysomes to 80S monosomes (P/M), measured in cycloheximide-stabilized extracts resolved by sedimentation through sucrose gradients (Fig. 5A-F), suggesting a reduced rate of bulk translation initiation relative to elongation in the mutants. No perturbation to the ratio of free 40S to free 60S subunits was evident in these gradients except for a modest reduction in the *R116D* mutant (Fig. 5A-F); even in this mutant, however, there was no significant decrease in the ratio of bulk 40S to 60S subunits in extracts prepared without cycloheximide and magnesium wherein polysomes and 80S monosomes dissociate into subunits (Fig. S1). These findings suggest that the alterations in accuracy of start codon selection observed in these mutants arise from altered 40S function, and not from abnormalities in expression of Rps3, 40S biogenesis, or stability of mature 40S subunits. This is consistent with the fact that scanning occurs only after assembly and attachment of 43S PICs to the mRNA, and so the fidelity of start-codon recognition during scanning should not be influenced by the concentration of 43S PICs.

### Rps3 Ssu^-^ substitutions R116D and R117D destabilize the P_IN_ conformation of the 48S PIC at UUG codons in vitro

The multiple defects in start codon recognition conferred by the *RPS3* mutations suggest that they might destabilize the P_IN_ state of the 48S PIC at both UUG and AUG start codons. We tested this hypothesis by analyzing their effects on the rate of TC dissociation from PICs reconstituted in vitro. To this end, we purified 40S subunits from *rps3Δ::kanMX* deletion strains harboring either plasmid-borne *R116D, R117D*, or WT *RPS3* as the only source of Rps3. 48S PICs were formed by incubating WT TC (assembled with [^35^S]-Met-tRNA_i_^Met^ and non-hydrolyzable GTP analog GDPNP) with saturating amounts of eIF1, eIF1A, an uncapped unstructured model mRNA containing either an AUG or UUG start codon [mRNA(AUG) or mRNA(UUG)], and either WT or mutant 40S subunits. 48S PICs containing [^35^S]-Met-tRNA_i_^Met^ were chased with excess unlabeled TC, incubated for increasing time periods, and then resolved via native gel electrophoresis to separate 40S-bound and unbound fractions of TC. In agreement with previous findings (Kolitz et al. 2009; Dong et al. 2014; Martin-Marcos et al. 2014), little to no TC dissociation occurred from WT PICs formed with mRNA(AUG) over the time course of the experiment (Fig. 6A-B), whereas appreciable dissociation was observed from WT PICs assembled on UUG-containing mRNA (k_off_ = 0.19±0.07 h^-1^; Fig. 6A-B). Neither Rps3 substitution substantially alters the kinetics of TC dissociation from PICs assembled on mRNA(AUG), although R117D generally conferred a moderate increase in the extent of dissociation (Fig. 6A-B, AUG mRNAs). By contrast, both the extent and rate of TC dissociation were substantially increased for PICs assembled on mRNA(UUG) using either R116D or R117D mutant 40S subunits compared to WT subunits (Fig. 6A-B). Previous work suggests that it is the P_IN_ conformation, and not the less stable P_OUT_ conformation, which is detected by these experiments and from which TC dissociation occurs. The rate of dissociation thus reflects the stability of the P_IN_ conformation, whereas the extent of dissociation reflects the ratio of PICs in P_IN_ versus a distinct, hyper-stable conformation from which TC does not detectably dissociate (Kolitz et al. 2009; Dong et al. 2014). Thus, the results in Fig. 6 indicate that Rps3 substitutions R116D and R117D substantially diminish rearrangement to the hyper-stable state and also markedly destabilize the P_IN_ state at near-cognate UUG codons.

**Figure 6.**
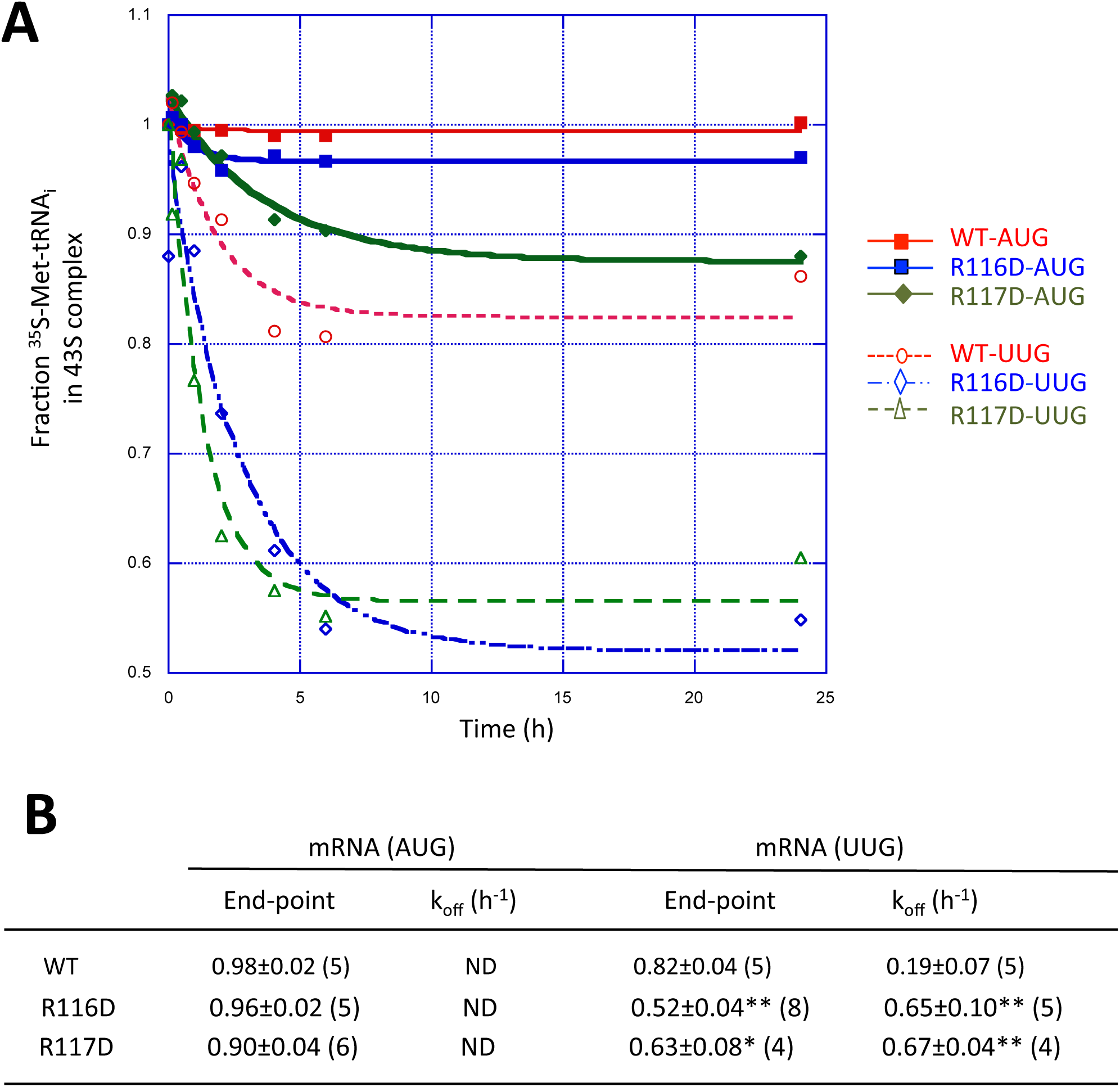
Rps3 Ssu^-^ substitutions R116D and R117D destabilize the P_IN_ conformation of the 48S PIC at UUG codons in vitro. (**A**) Analysis of TC dissociation from 43S-mRNA complexes assembled with WT, Rps3-R116D, or Rps3-R117D 40S subunits and either mRNA(AUG) or mRNA(UUG). Representative curves are shown for each measurement. (**B**) The endpoints and K_off_ values (±S.E.M.s) determined from between four and nine replicate experiments (number in parenthesis); ND, dissociation was too limited in most replicate determinations to permit k_off_ calculations. Asterisks indicate significant differences between mutant and WT as judged by a two-tailed, unpaired Student’s t-test (*, P<0.05; **, P<0.01).

### Rps3 Ssu^-^ substitutions R116D and R117D destabilize mRNA binding at the 40S entry channel

We recently analyzed the role of eIF3 subunits and domains in mediating stable interactions of the PIC with mRNA nucleotides located at the entry or exit channel of the 40S subunit by reconstituting PICs with different model mRNAs designed to incompletely occupy either the entry or exit channel of the 40S subunit (Aitken et al. 2016). Capped model mRNAs designated “5’5-AUG” and “5’11-AUG” contain only 5 or 11 nt located 5’ of the AUG start codon, respectively, but >30 nt 3’ of the AUG. Hence, with the AUG positioned in the 40S P site, both mRNAs should fully occupy the entry channel and protrude from the entry channel opening on the solvent-exposed surface of the 40S subunit, but contain only 2 or 8 nt in the exit channel beyond the 3 nt occupying the E-site (positions -3, -2, and -1 relative to the AUG; Fig. 7A). Because the exit channel accommodates ~10 nt, it is largely empty for PICS assembled on 5’5-AUG mRNA, but nearly filled for those assembled on 5’11-AUG mRNA; although neither mRNA will protrude outside the exit channel pore. Conversely, model mRNAs “3’5-AUG” and “3’11-AUG” contain >30 nt upstream of the AUG codon, and thus should fully occupy the exit channel, but will contain only 2 or 8 nt in the entry channel in addition to the 3 nt that occupy the A site (positions +4 to +6) when the AUG codon is in the 40S P site (Fig. 7A). Thus, the ~9 nt-long entry channel should be largely empty in PICs assembled on 3’5-AUG but fully occupied in PICs assembled on 3’11-AUG mRNA; neither mRNA protrudes beyond the entry channel pore. As a positive control, we also examined “mid-AUG” mRNA, containing residues both 5’ and 3’ of the AUG sufficient to fully occupy both the entry and exit channels and extend outside the openings of both channels (Fig. 7A). Radiolabeled versions of these mRNAs were employed to assay 48S PIC assembly in the reconstituted system, at saturating concentrations of 40S subunits, TC (assembled with GDPNP), eIF1, eIF1A, eIF5, eIF4A, eIF4B, the eIF4E•eIF4G complex, and eIF3, using native gel electrophoresis to separate 43S PIC-bound mRNA from unbound mRNA. While the eIF4 group of factors are dispensable for 48S PIC assembly using the uncapped model mRNAs employed for the TC dissociation assays in Fig. 6, they were included here to insure rapid and complete recruitment of the capped mRNAs employed here (Aitken et al. 2016).

**Figure 7.**
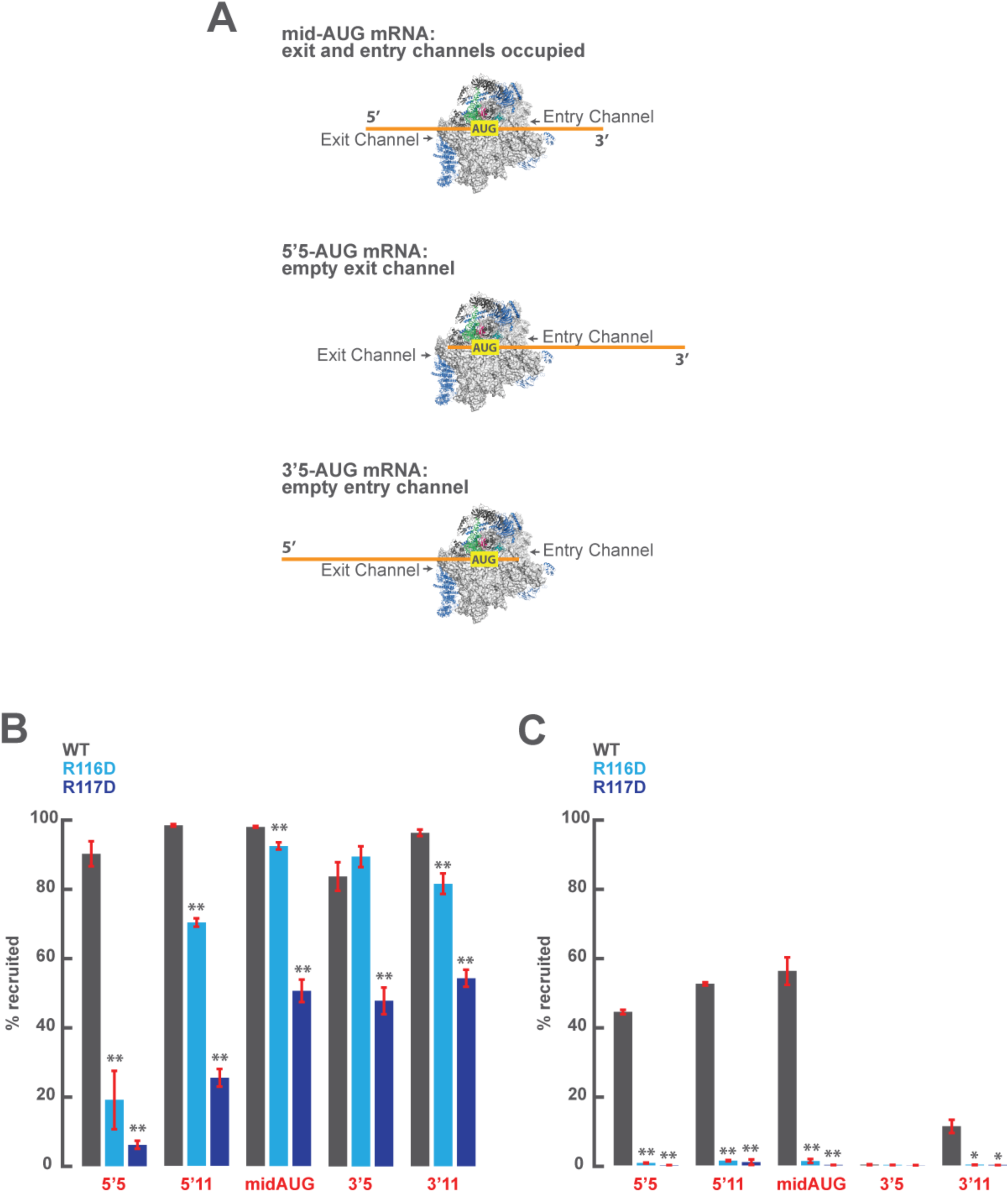
Rps3 Ssu^-^ substitutions R116D and R117D destabilize PIC:mRNA interaction at the 40S entry channel. (**A**) Schematic representation of 48S PICs formed in the presence of the mid-AUG, 5’5-AUG, and 3’5-AUG mRNAs. While the mid-AUG mRNA programs a complex in which both the mRNA entry and exit channels are occupied, the 5’5-AUG and 3’5-AUG mRNAs program complexes that leave either the exit or entry channels largely unoccupied, respectively. (**B**) The maximal extent of recruitment observed for capped model mRNAs in the presence of all factors, including eIF3, to PICs assembled with 40S subunits containing WT (grey), R116D (light blue), or R117D (dark blue) Rps3. (**C**) The maximal extent of recruitment observed for model mRNAs in the presence of all factors, with the exception of eIF3, to PICs assembled with 40S subunits containing WT (grey), R116D (light blue), or R117D (dark blue) Rps3. Asterisks indicate significant differences between mutant and WT as judged by a two-tailed, unpaired Student’s t-test (*, P<0.05; **, P<0.01).

Consistent with our previous study, 80-100% of each of these mRNAs can be driven into 48S PICs when eIF3 is present together with all other components and WT 40S subunits (Fig. 7B, grey bars), whereas omitting eIF3 reduces the extent of recruitment of the 5’5-AUG, 5’11-AUG, and mid-AUG mRNAs by ~40-60%, drastically impairs recruitment of 3’11 mRNA, and abolishes recruitment of 3’5 mRNA (Fig. 7C, grey bars). Similar results previously led us to conclude that eIF3 mediates interactions of the PIC with mRNA residues at the exit channel, which become crucial for mRNA recruitment when the entry channel is largely empty (as for 3’5-AUG mRNA) or when no mRNA residues protrude from the entry channel pore (for 3’11-AUG mRNA). eIF3 also enhances PIC·mRNA interactions at the entry channel, increasing recruitment of mRNAs that fully occupy the entry channel even when eIF3·mRNA interactions at the exit channel are impaired, as with 5’5 mRNA (cf. in Figs. 3B-C, 5’5 mRNA, grey bars) (Aitken et al. 2016). The fact that eIF3 is essential for mRNA recruitment only when mRNA nucleotides are lacking at the entry channel (as for 3’5 mRNA; Fig. 3C) led us to hypothesize here that contacts between Rps3 and mRNA at the entry channel promote mRNA recruitment by the PIC independently of eIF3. Because this compensatory interaction would be lost or impaired for the 3’5 and 3’11 mRNAs lacking mRNA residues at the entry channel, this would explain why eIF3 interactions with mRNA at the exit channel are essential for recruitment of these mRNAs. Indeed, the Rps3 residues studied here are located in proximity to mRNA nucleotides both within and just outside of the entry channel opening in the cryo-EM structure of py48S (Hussain et al. 2014b) (Fig. 1C).

Remarkably, when WT 40S subunits were replaced with R116D mutant subunits in reactions containing eIF3, we observed a dramatic reduction in recruitment of the 5’5 mRNA, which leaves the exit channel largely empty (Fig. 7B, cyan bars). This indicates that Arg-116 in Rps3 is crucial for PIC-mRNA interactions at the entry channel, and that these interactions are essential for mRNA recruitment when PIC-mRNA interactions at the opposite, exit channel are missing. Supporting this interpretation, the magnitude of the defect using R116D subunits is progressively diminished when the exit channel is filled by using 5’11-AUG mRNA and then when mRNA protrudes from the exit channel pore by using mid-AUG mRNA. The defect is also eliminated entirely using the 3’5-AUG or 3’11-AUG mRNAs (Fig. 7B, cyan bars), which lack mRNA bases at the entry channel but fully occupy and protrude from the exit channel. These last findings are consistent with our previous observation that interactions at the entry channel are redundant with those at the exit channel (Aitken et al. 2016), and also suggest that R116D does not impair other aspects of 40S structure or function that would compromise stable 48S PIC assembly with these mRNAs beyond the loss of mRNA·40S interactions at the entry channel.

Similar results were obtained when WT 40S subunits were replaced with R117D mutant subunits in reactions containing eIF3, including the nearly complete loss of 5’5 mRNA recruitment, which is partially rescued with 5’11 mRNA and more completely rescued with mid-AUG, 3’5, and 3’11 mRNAs (Fig. 7B, blue bars). In contrast to the results for R116D, however, the latter three mRNAs do not fully rescue recruitment to the level observed with WT or Rps3-R116D 40S subunits (Fig. 7B, blue vs cyan/grey bars). This last distinction might indicate that the R117D substitution impairs another aspect of stable 48S PIC assembly in addition to mRNA interactions at the entry channel.

Finally, in reactions lacking eIF3, we found that both Rps3 substitutions essentially abolished the appreciable mRNA recruitment retained in the absence of eIF3 for the mRNAs that completely occupy the entry channel (5’5, 5’11, and mid-AUG, Fig. 7C, blue bars). These findings fully support our conclusion that Rps3·mRNA interactions involving R116/R117 are essential for the eIF3-independent interaction of the PIC with mRNA at the entry channel. These results further provide strong evidence that direct contacts of Rps3 residues R116/R117 with mRNA nucleotides at the entry channel cooperate with eIF3 interactions at both the exit and entry channels to stabilize mRNA binding to 43S PICs.

## DISCUSSION

In this study, we obtained genetic and biochemical evidence implicating Rps3 residues at the 40S mRNA entry channel in promoting bulk translation initiation and maintaining wild-type accuracy of start codon recognition in vivo, in stabilizing the P_IN_ conformation of TC binding to the PIC, and in stabilizing interactions of the PIC with mRNA nucleotides at the entry channel that complement eIF3-mediated PIC:mRNA interactions at the entry and exit channels. In the recent py48S cryo-EM structure (Hussain et al. 2014a), mRNA in the entry channel was visualized up to residue +12, where it emerges from the entry channel pore on the solvent-exposed surface of the 40S subunit, with nucleotides +8 to +12 in proximity to basic residues of Rps3 (Fig. 1C). We found that Asp substitutions of three such residues, R116, R117, and R146, and the nearby basic residue K108, moderately reduced the rate of bulk translation initiation, as indicated by a decreased ratio of polysomes to monosomes, without decreasing the ratio of bulk 40S to 60S subunit abundance. These substitutions dramatically suppressed the increased utilization of UUG start codons in *HIS4* mRNA engendered by the Sui^-^ variant of eIF5 encoded by *SUI5*, thus conferring the Ssu^-^ phenotype. The R116, R117, R146, and K148 substitutions also reduced expression of eIF1, an effect observed previously for Ssu^-^ mutations of eIF1 and eIF1A and attributed to increased discrimination against the poor Kozak context of the eIF1 start codon (Martin-Marcos et al. 2011). We established that this mechanism also applies to the Rps3 R116D and R117D substitutions by showing that they reduce expression of a *SUI1-lacZ* fusion containing the native, poor context for the (eIF1) AUG start codon, but not that of a modified *SUI1-lacZ* reporter containing optimum Kozak context. It is unclear however whether this mechanism underlies the relatively weaker reductions in eIF1 levels conferred by the K148D and R146D substitutions. Thus, R116D and R117D increase discrimination against the eIF1 AUG start codon in its native, poor context as well as reducing recognition of near-cognate UUG codons, whereas K148D and R146D only strongly suppress UUG initiation. Our previous genetic analyses of residues of the β-hairPin of Rps5, which is located in the 40S mRNA exit channel, provides a precedent for altered 40S contacts with mRNA that increase discrimination against a mismatch in the start-codon:anticodon duplex at UUG codons without altering the influence of Kozak context (Visweswaraiah et al. 2015). As discussed further below, this might indicate that these two aspects of start codon recognition have distinct molecular mechanisms.

As R116D and R117D conferred the broadest and most pronounced genetic defects, we purified mutant 40S subunits harboring these Rps3 substitutions and analyzed their effects on the stability of either TC or mRNA binding to the PIC in the yeast reconstituted system. We found that both R116D and R117D substantially increased the dissociation rate of TC from PICs reconstituted with a model unstructured mRNA containing a UUG start codon, without significantly affecting the corresponding AUG complex. These findings support the possibility that the suppression of UUG initiation conferred by these mutations in vivo (their Ssu^-^ phenotype) arises from destabilization of the P_IN_ state of TC binding in the presence of the inherently less stable codon:anticodon duplex containing a U:U mismatch that is formed at UUG codons. The discrimination against the *SUI1* AUG in native, poor Kozak context engendered by the R116D/R117D substitutions cannot be explained by the same mechanism, as the model mRNAs employed for k_off_ measurements contain poor Kozak context for the AUG/UUG start codons. Instead, perhaps R116D/R117D reduce recognition of the *SUI1* AUG in poor context by decreasing its dwell-time in the P site during scanning, an important factor in determining the impact of poor context on initiation in the mammalian system (Kozak 1990). If so, this effect might also contribute to the reduced recognition of UUG codons conferred by these Rps3 substitutions.

Using a complementary assay that employs model mRNAs that leave empty or only partially occupied the entry or exit channel of the 40S subunit within 48S PICS, we observed that both R116D and R117D dramatically destabilize mRNA interactions at the entry channel. We showed previously that eIF3 is crucial for stabilizing mRNA:PIC interactions at the exit channel, whereas substantial mRNA interactions at the entry channel can occur independently of eIF3 (Aitken et al. 2016). Here we found that the Rps3 R116D and R117D substitutions strongly destabilize 48S PICs assembled in the presence of eIF3 on 5’-truncated mRNAs lacking some or all of the eIF3-sensitive mRNA contacts in the exit channel (5’5-AUG and 5’11-AUG mRNAs). In the absence of eIF3, these Rps3 substitutions destabilize binding of PICs to all mRNA substrates except the one nearly devoid of mRNA in the entry channel (3’5 mRNA). However, this last mRNA does not bind stably to the PIC in the absence of eIF3 even when using WT 40S subunits, because it lacks contacts with the PIC at the entry channel, which we can now attribute to Rps3 residues Arg116 and Arg117.

Previous studies have shown that elongating bacterial 70S ribosomes exhibit a helicase activity, driven by mRNA translocation through the ribosome, that enables efficient elongation through highly structured mRNA sequences, and that basic residues in uS3 and uS4 at the opening of the mRNA entry channel are required for the helicase activity in vitro, including R131, R132, and K135 in uS3 (Takyar et al. 2005). Remarkably, alignment of uS3 sequences from eukaryotic, archaebacterial, and bacterial sources indicates that residues R131 and R132 in bacterial uS3 align with R116 and R117 in yeast uS3 (Graifer et al. 2014). It has been proposed that the conserved basic residues in uS3 and uS4 contribute to ribosome helicase function by interacting with the phosphate backbone of the complementary strands of an mRNA duplex positioned at the entry channel pore; these interactions would either stabilize the unwound strands generated by spontaneous melting of the duplex or function more directly to couple 30S head movement during translocation to mRNA duplex unwinding (Takyar et al. 2005). Because the model mRNAs used in our experiments were designed to be devoid of secondary structure, it seems unlikely that Rps3 residues R116/R117 promote PIC-mRNA interactions at the entry channel by helping to melt secondary structures in the mRNA. Instead, they might simply provide electrostatic attraction for the mRNA phosphate backbone, helPINg to fix mRNA to the head of the 40S subunit, where Rps3 resides. In this view, basic residues in Rps30/eS30, also located at the entry channel opening, could similarly fix the mRNA to the 40S body. All of these interactions would occur simultaneously when the head moves closer to the body as a result of the transition from the open to the closed conformation of the PIC that is provoked by AUG recognition (Llacer et al. 2015) (see model in Fig. 8). These P_IN_cers would thus help to clamp the mRNA into the now constricted entry channel, arresting scanning and helping to stabilize the closed conformation, in which the P site is fully formed and encloses the Met-tRNA_i_^Met^ in the P_IN_ state. Thus, by eliminating mRNA interactions at the entry channel, the Rps3 R116D/R117D substitutions would destabilize the closed conformation of the PIC and indirectly destabilize Met-tRNA_i_^Met^ binding in the P site. Combining this destabilizing effect with the less stable codon:anticodon duplex formed at UUG codons could account for the increased rate of TC dissociation from PICs reconstituted with R116D/R117D mutant 40S subunits and mRNA(UUG) (Fig. 6), as well as the increased discrimination against UUG codons (Ssu^-^ phenotype) conferred by these substitutions in vivo (Fig. 3).

**Figure 8.**
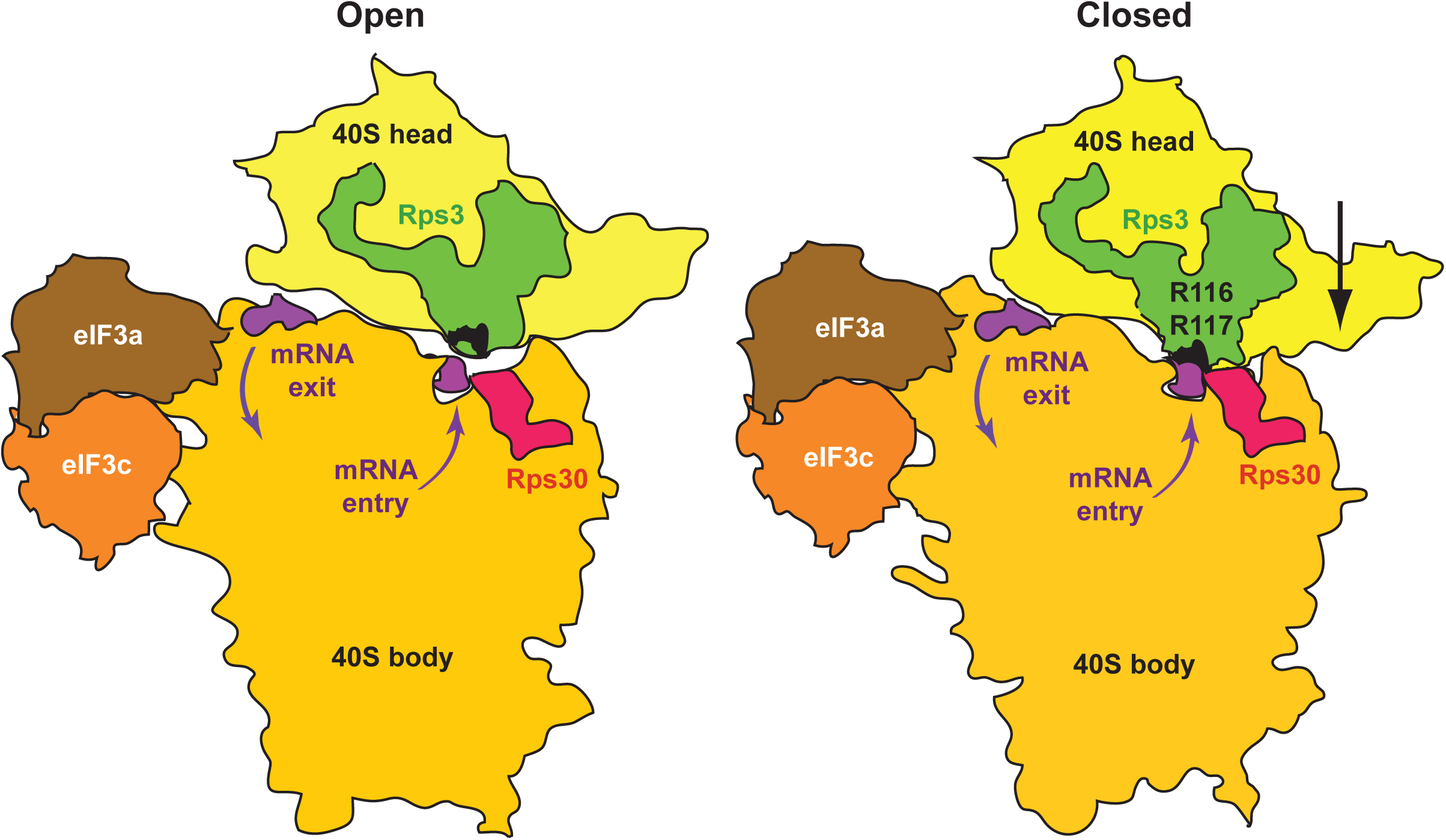
Model depicting a conformational switch in the PIC between the open, scanning and closed conformations on start codon recognition that clamps mRNA at the entry channel between Rps3 and Rps30. Silhouettes of 40S subunits were traced from py48S-open (PDB: 3jaq) and py48S (PDB: 3j81) to depict the open and closed conformations of the PIC, respectively. PCI domains of eIF3 a/c subunits were traced from py48S-closed (PDB 3jap), and mRNA (purple) was traced from py48S (PDB 3j81), and docked at similar positions on both 40S silhouettes. Locations of R116/R117 are shown in black on the green silhouettes of Rps3. Downward movement of the 40S head towards the body in the closed complex relative to its position in the open complex (Llacer et al. 2015) (depicted by arrow on the right) narrows the mRNA entry channel, engaging R116/R117 with mRNA at the entry channel pore. These contacts, and interactions of Rps30 (red) with mRNA at the same location help fix the mRNA in the entry channel and stabilize the closed/P_IN_ state at the start codon. The eIF3a PCI domain stabilizes interaction of the PIC with mRNA at the exit channel (Aitken et al. 2016), complementing the function of Rps3 R116/R117 at the entry channel.

Finally, it is interesting that Rps3 amino acids Arg-146 and Lys-141 also interact with rRNA residues in helices 18 and 34 that are involved in the non-covalent interactions between rRNA residues in the 40S body and head. These interactions compose the latch over the mRNA entry channel, with Arg-146 interacting with h34 in both the open and closed conformations and Lys-141 interacting with both helices but with h18 only in the closed state (Llacer et al. 2015). These Rps3:rRNA interactions should promote the closed conformation of the latch, which in turn should help to clamp mRNA in the exit channel and stabilize the closed arrangement of the 40S head relative to the body that locks Met-tRNA_i_^Met^ into the P site. Thus, the diminished UUG initiation we observed for an acidic substitution of R146 might involve a weakened entry channel latch in addition to diminished Rps3:mRNA contacts in the entry channel. The strong growth defect observed for the K141D substitution is consistent with an even greater destabilization of the closed conformation of the PIC resulting from a destabilized latch. Given the collaboration of Rps3 with eIF3 in stabilizing mRNA at the entry channel, it will be interesting to learn whether the conserved basic residues in Rps3 functionally interact with segments of eIF3 subunits, in particular the eIF3a CTD, which has been implicated in PIC:mRNA interactions at the entry channel (Chiu et al. 2010; Aitken et al. 2016) and in start codon selection (Valasek 2012).

## MATERIALS AND METHODS

### Plasmid constructions

Plasmids employed in this work are listed in Table S3. Plasmid pDH412 was constructed as follows. The *RPS3* gene, including 458 bp upstream of the ATG start codon and 200 bp downstream of the stop codon, was amplified by PCR using the two primers 5′-GAA TGC GGC CGC GAA GCA GTT ACA TCT CA-3′ and 5′AAG TCG ACT TGT GTT GTA CAA AAC TT-3′ and genomic DNA from yeast WT strain BY4741 as template. The ~1.4 kb amplicon was digested with NotI and SalI, and the resulting fragment was inserted between the NotI and SalI sites of plasmid pRS315 to produce pDH412. The DNA sequence of the entire open reading frame (ORF) was verified. pDH459 was constructed by inserting the NotI-SalI fragment containing *RPS3* isolated from pDH412 between the NotI and SalI sites of pRS316. It was shown that both pDH412 and pDH459 fully rescue growth of *P_GAL_-RPS3* strain HD2738 on glucose medium. Plasmids pDH424, pDH431, pDH425, pDH432, pDH482, pDH433, pDH13-37, pDH429, and pDH430 were derived from pDH412 by site-directed mutagenesis using the QuickChange XL kit (Stratagene) and the corresponding primers in Table S4, and the mutations were verified by sequencing the entire ORF and 5’ non-coding region.

### Yeast strain constructions

Yeast strains employed in this work are listed in Table 1.

Strain HD2738 was derived from H2995 by replacing the promoter of chromosomal *RPS3* with the *GAL1* promoter by one-step gene replacement, as follows. Primers (i) 5’-CCT TTC CTG TAT AAT ATT CTT GCT GTA AAG TTT GTT TTT TTT ATG AAA AAA ACA TTT TCT TTT CTT GAG GAA TTC GAG CTC GTT TAA AC and (ii) 5’ GAA TTC GTT CAA TTC AGC GTA GAA GAC ACC GTC AGC GAC TAG CTT TCT TTT CTT AGA GAT TAA AGC GAC CAT TTT GAG ATC CGG GTT TT were used to amplify by PCR the appropriate DNA fragment from plasmid pFA6a-kanMX6-PGAL1 (p3218), which was used to transform strain H2995 to kanamycin resistance. The presence of *P_GAL1_-RPS3* in the resulting strain (HD2738) was established by demonstrating that the lethality on glucose medium can be complemented by lc *RPS3 LEU2* plasmid pDH412, and confirmed by PCR analysis of chromosomal DNA with the appropriate primers.

Strains HD2754, HD2755, HD2765, HD2764, HD2767, HD2766, HD2769, HD2768, HD3120, HD2772, and HD2779 were derived from HD2738 by transformation with a *LEU2* plasmid containing the indicated *RPS3* allele, or empty vector, as indicated in Table S2. Strains HD2836, HD2911, HD2846, HD2868, HD2848, HD2861, HD2850, HD2849, HD3145, HD2841, and HD2860 were derived from the foregoing strains by transformation with sc *TRP1 SUI5* plasmid p4281), or empty vector YCplac22, as indicated in Table S2.

To produce strain HD2973, pDH459 (lc *URA3 RPS3*^+^) was introduced into the *RPS3/rps3*Δ diploid strain YSC1021-671817 purchased from Research Genetics. The Ura^+^ transformants were sporulated and subjected to tetrad analysis. HD2973 was identified as an Kan^R^ Ura^+^ ascospore clone incapable of growth on medium containing 5-FOA unless transformed with *RPS3+ LEU2* plasmid pDH412 but not with empty *LEU2* vector pRS315. The presence of *rps3Δ::kanMX* in HD2973 was verified by PCR analysis of chromosomal DNA. The strains HD3240, HD3241, HD3242, HD3243, and HD3244 were derived from HD2973 by plasmid shuffling to replace pDH459 with the appropriate lc *LEU2* plasmid containing the *RPS3* allele indicated in Table S2.

### Biochemical analysis of yeast cells

Assays of β-galactosidase activity in whole-cell extracts (WCEs) were performed as described previously (Moehle and Hinnebusch 1991). For Western analysis, WCEs were prepared by trichloroacetic acid extraction as described (Reid and Schatz 1982), and immunoblot analysis was conducted as described previously (Martin-Marcos et al. 2011) with antibodies against eIF1 (Valasek et al. 2004) (Valášek et al. 2001). The signal intensities were quantified using a LI-COR Odyssey infra-red scanner.

For polysome analysis, strains HD2754 (*RPS3*^+^), HD2767 (*R116D*), HD2769 (*R117D*), HD3120 (*K108D*), HD2765 (*K148D*), or HD2767 (*R146D*) were grown in SC-Leu medium at 30°C to A_600_ ~1. Cycloheximide was added to 50 μg/ml 5 min prior to harvesting, and WCE was prepared in breaking buffer (20 mM Tris–HCl, pH 7.5, 50 mM KCl, 10 mM MgCl_2_, 1 mM dithiothreitol, 5 mM NaF, 1 mM phenylmethylsulfonyl fluoride, 1 Complete EDTA-free Protease Inhibitor Tablet (Roche)/50 ml buffer). 15 A_260_ units of WCE were separated by velocity sedimentation on a 4.5–45% sucrose gradient by centrifugation at 39,000 rpm for 3 h in an SW41Ti rotor (Beckman). Gradient fractions were scanned at 254 nm to visualize ribosomal species. To analyze total 40S/60S profiles, the *rps3Δ::kanMX* deletion strains harboring plasmid-borne WT *RPS3*^+^ (HD3240) or *rps3* alleles *R116D* (HD3241), *R117D* (HD3242), *K108D* (HD3243), *K148D* (HD3244) or *R146D* (HD3277) were grown in YPD at 30°C to A_600_~1, but cycloheximide was omitted. WCEs were prepared in the absence of Mg^+2^ and 15 A_260_ units of WCE were resolved by velocity sedimentation through 5 to 30% sucrose gradients at 40,000 rpm for 4 h in an SW41Ti rotor (Beckman).

### Biochemical analysis in the reconstituted yeast translation system

#### TC dissociation rates

Initiation factors eIF1A, eIF1, and His6-tagged eIF2 were purified as described (Acker et al. 2007). WT and mutant 40S subunits were purified from the *rps3Δ::kanMX* deletion strains harboring plasmid-borne WT *RPS3*^+^ (HD3240), *rps3-R116D* (HD3241), or *rps3-R117D* (HD3242), as the only source of Rps3, as described previously (Acker et al. 2007). Model mRNAs with sequences 5′-GGAA[UC]_7_UAUG[CU]_10_C-3′ and 5′-GGAA[UC]_7_UUUG[CU]_10_C-3′ were purchased from Thermo Scientific. Yeast tRNA_i_^Met^ was synthesized from a hammerhead fusion template using T7 RNA polymerase, charged with [^35^S]-methionine, and used to prepare radiolabeled eIF2·GDPNP·[^35^S]-Met-tRNA_i_^Met^ ternary complexes ([^35^S]-TC), all as previously described (Acker et al. 2007). Yeast Met-tRNA_i_^Met^ was purchased from tRNA Probes, LLC and used to prepare unlabeled TC in the same way. Dissociation rates of TC (k_off_) were measured by monitoring the amount of [^35^S]-TC that remains bound to 40S·eIF1·eIF1A·mRNA complexes over time, in the presence of excess unlabeled TC (chase), using native gel electrophoresis to separate 40S-bound from unbound [^35^S]-TC. 43S·mRNA complexes were preassembled for 2 h at 26°C in reactions containing 40S subunits (20 nM), eIF1 (1 μM), eIF1A (1 μM), mRNA (10 μM), and [^35^S]-TC (0.25 μM eIF2/0.1 mM GDPNP/1 nM [^35^S]-Met-tRNA_i_^Met^) in 60 μl of 1X Recon buffer. To initiate each dissociation reaction, a 6 μl-aliquot of the preassembled 43S·mRNA complexes was mixed with 3 μl of 3-fold concentrated unlabeled TC chase (comprised of 2 μM eIF2/0.3 mM GDPNP/0.9 μM Met-tRNA_i_^Met^), to yield a 300-fold excess of unlabeled versus labeled TC in the final dissociation reaction, and incubated for the prescribed period of time. A converging time course was employed so that all dissociation reactions are terminated simultaneously by the addition of native-gel dye and loaded directly on a running native gel. The fraction of [^35^S]-Met-tRNA_i_ remaining in 43S complexes at each time point was determined by quantifying the 40S-bound and unbound signals by phosphorimaging, normalized to the ratio observed at the earliest time-point, and the data were fit with a single exponential equation (Kolitz et al. 2009).

#### mRNA recruitment to 43S PICs

Preparation of initiation factors, charged Met-tRNA_i_^Met^, and 40S ribosomal subunits were purified as previously described (Acker et al. 2007; Mitchell et al. 2010; Aitken et al. 2016). Model mRNAs described by Aitken et al (2016) were transcribed in vitro, capped with [α-^32^P]-GTP (PerkinElmer) using vaccinia virus D1/D12 capping enzyme, and purified as described (Mitchell et al. 2010), with the modifications described by Aitken et al (2016). The extent of mRNA recruitment to reconstituted 43S PICs was determined using a previously-described gel-shift assay (Mitchell et al. 2010). PICs were assembled in the presence of 1 μM eIF1, 1 μM eIF1A, 300 nM eIF2, 200 nM Met-tRNA_i_^Met^, 400 nM eIF3, 2 μM eIF4A, 300 nM eIF4B, 50 nM eIF4E•eIFG, 300 nM eIF5, and 30 nM 40S subunits in 1X Recon buffer (30 mM HEPES-KOH pH 7.4, 100 mM KOAc pH 7.6, 3 mM Mg(OAc)2, 2 mM DTT) and incubated 10 min at 26 °C. Reactions were initiated by the simultaneous addition of ATP•Mg^2+^ and the appropriate [^32^P]-capped mRNA to final concentrations of 2 mM and 15 nM, respectively, and incubated for 2 h at 26 °C, at which point all reactions had proceeded to completion as judged by prior kinetic experiments (Aitken et al. 2016). The free and PIC-bound mRNA fractions were resolved on a 4% native THEM gel, run for 45 min at 200 V, and quantified by phosphorimaging analysis using a Typhoon FLA 57 9500 imager (GE Life Sciences). The extent of recruitment was calculated as the fraction of total signal in the lane represented by PIC-bound mRNA.

## ACKNOWLEDGEMENTS

We thank Jagpreet Nanda, Jyothsna Visweswaraiah, and Fan Zhang for advice on k_off_ determinations. We are grateful to members of our laboratories and Tom Dever’s group for helpful suggestions. This work was supported in part by the Intramural Research Program of the National Institutes of Health (A.G.H. and J.R.L.) and NIH grant GM62128 (previously to J.R.L.). C.E.A. was further supported by an NIH minority supplement to NIH grant GM62128 and by a Leukemia & Lymphoma Society CDP Fellowship.

